# Encoding and Decoding of Brain Dynamic Functional Connectivity for ADHD Diagnosis

**DOI:** 10.64898/2025.12.05.692480

**Authors:** Deepank Girish, Yi Hao Chan, Sukrit Gupta, Jing Xia, Jagath C. Rajapakse

## Abstract

Recent studies have demonstrated strong associations between the changes in dynamic functional connectivity (FC) and both behavioral and cognitive functions. The sliding window technique is the most widely used method for evaluating dynamic FC; however, it faces two key challenges: distributional shifts across windows and high dimensionality, as FC is computed across windows of the entire time series. To address these issues, we propose BRAINMAP (Bi-level Representation using Attention for INterpretability with Mamba-Aided Prediction) to model the dynamic FC of the brain. BRAINMAP employs the Optimal Transport technique to correct distributional shifts across sliding windows and leverages Graph Neural Networks (GNNs) in conjunction with a hybrid approach that integrates an attention mechanism and the Mamba block to effectively capture spatiotemporal features for functional MR images. Finally, we introduce a novel Top-K sliding window feature selection algorithm to induce sparsity in dynamic FC. We conducted an extensive evaluation of our model for diagnosing Attention Deficit Hyperactivity Disorder (ADHD) using three resting-state fMRI datasets: ADHD-200, UCLA, and CNI-TLC, which comprise a total of 447 subjects with ADHD and 845 typically developing controls. Our architecture outperformed existing state-of-the-art dynamic FC models in ADHD detection, achieving improvements ranging from 3% to 12% across the three datasets. We demonstrate that our proposed model produces robust biomarkers, most notably the connection between the dorsal attention network and the visual network. Using an association study, we further establish the clinical relevance of the identified biomarkers.

## I Introduction

**A**MONG the most widely used neuroimaging modalities, functional magnetic resonance imaging (fMRI) stands out as a powerful, noninvasive technique for measuring brain activity. It has been instrumental in advancing our understanding of a wide range of brain states, including those associated with neurodevelopmental conditions. Traditional fMRI analysis focused on static functional connectivity (FC) [1], [2], typically defined as the correlation between spatially distinct brain regions for the entire duration of the scan. Dynamic FC extends traditional FC by capturing temporal variations in brain-wide connectivity patterns as they evolve over time [3]. Recent converging evidence suggests that dynamic FC analysis is more effective in detecting neurodevelopmental disorders compared to static FC methods [4]. This is particularly relevant to ADHD, for which dynamic brain changes may better reflect underlying developmental delays [5].

To estimate dynamic FC, a widely adopted approach is the sliding window technique [6], where the BOLD time series is segmented into overlapping temporal windows and FC is computed within each segment. Although effective in capturing temporal transitions, this method overlooks the temporal covariate shift (distribution shift) across sliding windows [7]. Each window may potentially capture a different statistical representation of the underlying neural dynamics, leading to a non-stationary distribution of FC matrices. This variability hinders reliable disease classification by complicating the learning of dynamic brain patterns. Another issue to consider is that this approach introduces high dimensionality in dynamic FC analyses, which is further exacerbated by the limited sample size, as conventional models often face serious overfitting problems [8]. It is imperative to devise appropriate sparsity strategies for dynamic FC classification models.

To model such time-varying brain dynamics, graph neural networks (GNN) have emerged as a popular computational framework. Given the graph-structured nature of brain connectivity, GNNs have gained significant traction for modeling FC networks. GNNs have proven effective in classifying various neurodevelopmental disorders within dynamic FC settings [9]. One limitation of existing GNN-based approaches for FC network analysis is their inability to effectively capture temporal dynamics. Moreover, these models often lack temporal explainability. Addressing this limitation, attention mechanisms [10] have been integrated into dynamic FC frameworks to capture the inherent temporal features, thereby improving both the model’s classification performance and interpretability [11], [12]. However, attention mechanisms face two significant drawbacks: their restricted capacity to model dependencies beyond a fixed context window and quadratic time complexity with respect to sequence length.

To address these issues, we propose Bi-level Representation using Attention for INterpretability with Mamba-Aided Prediction (BRAINMAP) to model the brain’s functional architecture using dynamic FC. BRAINMAP addresses distributional shift by leveraging Optimal Transport, while spatial dependencies are captured using GNNs. We conjecture that integrating these two complementary representations enhances the model’s generalization capability, leading to improved classification performance. We subsequently employ Cross-Attention to combine the two representations and capture short-to medium-range temporal features. Additionally, the resulting attention scores are utilized for clinical interpretability, enabling the identification of salient features that contribute to the model’s predictions. To overcome the limitations of attention mechanisms, we incorporate Mamba [13] to effectively capture long-range dependencies across dynamic FC time windows. Mamba is a structured state space model (SSM), which has emerged as a robust sequence modeling architecture due to its principled approach to capturing longrange dependencies [14]. Furthermore, we propose a novel Top-K Sliding Window Selection algorithm that identifies the most informative dynamic FC windows, thereby mitigating overfitting and reducing computational overhead in parameter-intensive dynamic FC models.

Existing studies typically offer limited evaluation of their proposed biomarkers, often highlighting only the top-ranked features and relying on comparisons with prior studies that report similar results. Such evaluations usually overlook the possibility that the identified features may be misleading or spurious. To more robustly and objectively assess the salient features identified by our model, we employ a quantitative framework, RE-CONFIRM [15], which evaluates the reliability and robustness of the potential biomarkers highlighted. We then assess the clinical relevance of the identified biomarkers by examining their correlations with ADHD test scores.

The proposed model is extensively evaluated using resting-state fMRI datasets collected from individuals with ADHD and their controls, including the ADHD-200, UCLA, and CNI-TLC datasets. We demonstrate that our model consistently outperforms other baseline methods in detecting ADHD. Compared to the state-of-the-art techniques, our approach achieves a consistent 2% performance gains across all ADHD-200 sites (NYU, OHSU, and PKU), as well as on the UCLA and CNI-TLC datasets. We performed additional analyses to validate the effectiveness of our model, including biomarker analysis by using saliency scores from BRAINMAP’s predictions for ADHD classification across all three datasets.

The key contributions of this study are:

- We propose BRAINMAP, a framework for modeling brain dynamic FC by integrating the Cross-Attention mechanism with the Mamba module to handle long-term dependencies. Additionally, we address distributional shifts across sliding windows with an Optimal Transport (OT) mechanism.
- We develop a novel sliding window feature selection algorithm to address overfitting and reduce feature redundancy commonly encountered in dynamic FC classification tasks.
- By decoding the FC, we identified the dorsal attention–visual network connection as a salient feature for pediatric ADHD subjects. We also demonstrate how this connection correlates positively with ADHD symptom severity, as measured by Conners Scale scores.

## II. Methods

In this section, we outline the core components and design of BRAINMAP. The overall architecture of the proposed framework is illustrated in Figure 1. The input stage features two parallel branches that capture complementary representations: (i) OT models the global distribution shifts across sliding windows and (ii) GNN extracts spatial features. Next, a Cross-Attention layer is followed by a Mamba block, which combines the OT and GNN outputs to capture temporal dynamics. Finally, a novel Top-K Sliding Window Selection algorithm is employed to identify the most salient time windows for distinguishing between ADHD subjects and typically developing controls.

**Fig. 1.**
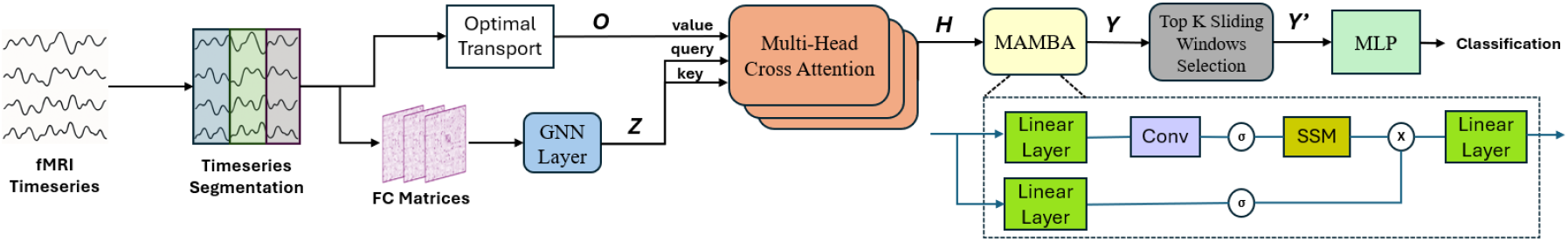
Overview of the proposed BRAINMAP architecture. Optimal Transport captures distributional shifts across sliding windows while GNNs extract spatial features. The outputs ***Z*** and ***O*** of GNN and OT, respectively, are fused via a multi-head Cross-Attention mechanism. The Mamba module then processes the combined representations ***H*** to capture temporal dependencies. Finally, the Top-K selection algorithm identifies the most informative windows for downstream prediction tasks.

### A. Graph Neural Networks

The brain FC is modeled as a graph *G*, where the set *V* of nodes corresponds to brain regions of interest (ROIs) and the set *E* of edges captures the connectivity between these ROIs. For each sliding window over time indices *t* ∈{1, 2, 3, …*T*}, a FC graph *G*_*t*_ = (*V, E*_*t*_, *X*_*t*_) is derived from resting-state fMRI (rs-fMRI) data. In this graph, the adjacency matrix *E*_*t*_ ∈ ℝ^*N ×N*^ encodes the strength of the connections between the ROIs, where *N* = |*V*| is the number of nodes in the set *V*. The matrix *X*_*t*_ ∈ ℝ^*N ×F*^ represents nodal features, with each feature defined by its Pearson correlation with all other nodes, and *F* indicates the dimensionality of the features of brain regions.

When constructing an adjacency matrix, researchers often aim to eliminate spurious connections likely caused by noise and to obtain a sparse FC network. This is achieved by selecting a predefined network density and computing a threshold that preserves the target edge density. However, there is no consensus in the literature regarding the optimal threshold [16], as weak correlations may still contain valuable functional information. To address this, we employed the orthogonal minimum spanning trees (OMST) filtering scheme [17] to threshold the adjacency matrix. This data-driven approach optimally balances the trade-off between minimizing noise by removing spurious edges and preserving beneficial weak connections.

GNN are extensively employed to capture both topological information, such as connectivity patterns, inter-regional relationships, and spatial information, including anatomical locations and geometrical properties of brain regions. Their effectiveness lies in their ability to leverage the complex information embedded within the graph’s structure and node attributes. GNNs are typically composed of two key functions: AGGREGATE and COMBINE. Different GNN variants are characterized by how they define and implement these functions. For example, GraphSAGE [18] uses neighborhood sampling with mean or LSTM-based aggregation. In contrast, GIN (Graph Isomorphism Network) [19] employs summation as the aggregation function and multi-layer perceptrons (MLPs) for combination. In most GNNs, for any node *v* ∈ *V* :

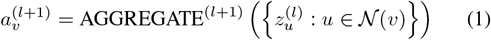

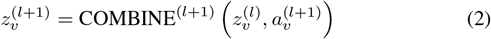

Here, *N* (*v*) represents the set of neighboring nodes of the central node *v*, and 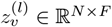denotes the feature vector of node *v* at layer *l*. The term 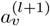 represents the aggregated information from the features of all neighboring nodes at layer *l* + 1. The COMBINE function updates the features of node *v* at layer *l* + 1 by applying a nonlinear transformation to the aggregated information 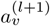 and the node’s own feature vector 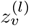 from the previous layer *l* + 1.

We evaluate the performance of various GNN architectures within our framework and ultimately employ a single-layer Graph Isomorphism Network (GIN) for spatial feature extraction. This layer takes the adjacency matrix *E*_*t*_ and the node feature matrix *X*_*t*_ as inputs. Node features are updated by aggregating information from neighboring nodes, producing a sequence of graph embeddings. The resulting sequence of graph embeddings 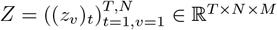, where *M* denotes the output dimension of the GNN, is then combined with the OT matrix and fed into the Cross-Attention layer.

### B. Distribution Shift Correction

In the sliding window method, given a predefined window length, FC is computed within each window, and the window is then shifted by one or more time points to repeat the estimation across the time series. A major limitation of the distributional shift in BOLD signals across time is often overlooked: When the input time series is divided into overlapping sliding windows, the underlying data distribution varies across these windows. These distributions are likely to differ from the actual distribution over the sample space, leading to covariate or distributional shifts driven by temporal non-stationarity. Shown in Figure 2, this results in poor generalization and thus hindering the performance and validity of dynamic FC models.

**Fig. 2.**
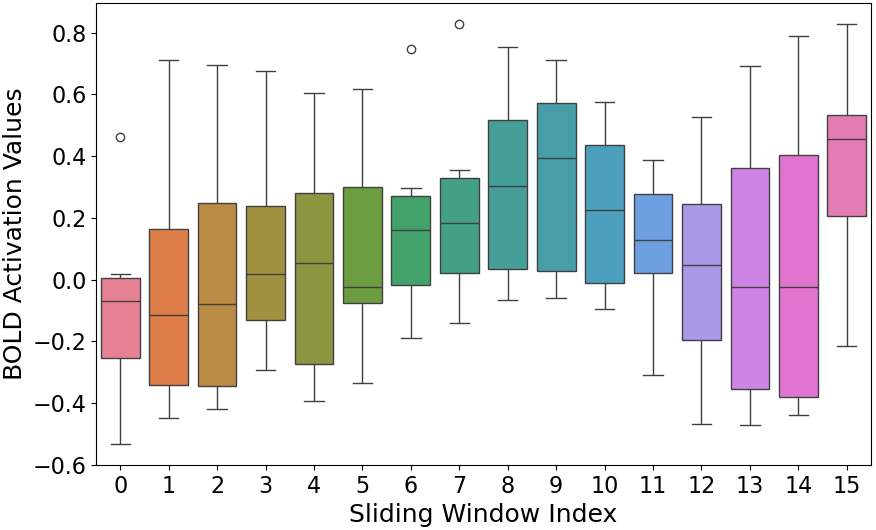
Illustration of distributional shift in dynamic FC across sliding windows. The bold activation values across moving windows were obtained by applying a sliding window of length 60 seconds and stride 16 seconds to the fMRI BOLD signals of brain regions parcellated by Power atlas, from site NI of ADHD-200.

To address the distribution shift, we utilize the Optimal Transport (OT) [20] mechanism that provides an optimal way to match or align two different distributions. OT first quantifies the difference between the source and target distributions by using a distance metric, such as the Wasserstein distance, which measures the “cost” of transforming one distribution to match the other. OT then seeks to find the “optimal” mapping (or transport map) by minimizing the cost of moving samples from the source to the target distribution.

We adopt the Kantorovich formulation [21] to compute the transport plan between the discrete distributions corresponding to each sliding window of fMRI time-series. The sliding window distributions across all brain regions are denoted as 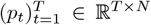. The objective is to compute the discrete transport plan matrix *O*_*t* −1_ ∈ ℝ^*N ×N*^, which maps the distribution from the *t* − 1^*th*^ sliding window to that of the *t*^*th*^ sliding window. Given ℒ is the cost of moving from *t* − 1^*th*^ to *t*^*th*^ distribution, the optimal transport plan (also referred to as optimal coupling) is given by the following optimization problem:

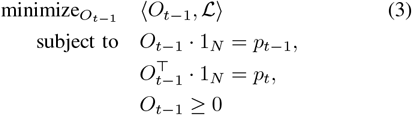

where ⟨.,. ⟩ denotes the Frobenius inner product of matrices and 1_*N*_ denotes a *N*-dimensional vector of ones. In the context of brain connectivity, ℒ represents the redistribution of a proportion of neural activity or connectivity strength from one region to another over time. As the network evolves, a portion of the connectivity remains within the same region while the rest is redistributed to other regions. The OT framework performs this redistribution while minimizing the overall cost, reflecting anatomical distance, functional similarity, or both. Such couplings (source and target window distributions) capture how brain activity redistributes over time and define distances in a latent space that reflect dynamic functional relationships.

In essence, OT not only mitigates distributional discrepancies but also captures temporal and structural relationships that better reflect underlying network dynamics. In the context of dynamic brain networks, this simulation models the global interaction among brain regions and illustrates the evolution of their topological structure. This was subsequently used as one of the inputs to the attention layer.

### C. Cross-Attention Layer

To integrate two feature representations obtained from OT and GNN, we utilize cross-attention modules. These modules utilize the multi-head attention (MHA) mechanism introduced in the Transformer architecture [10]. The output of each attention head, *H*_*i*_, in the multi-head attention module is expressed as follows:

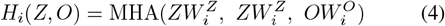

where *Z* denotes the embeddings from the GNN, and 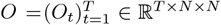 represents the embeddings obtained via OT. In our formulation, the graph embeddings serve as both the query and key, while the OT matrix is used for the value. As in the transformer formulation, 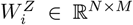 denotes the weight matrices used to compute the queries and keys, while 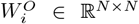 refers to the weight matrices used to compute the values in the attention mechanism of the *Z*^*th*^ head. The influence of the two sets of representations is evaluated sequentially for each sliding window, enabling the model to account for both local interactions and global differences in network organization. Finally, outputs from all attention heads are averaged, and the resulting features are concatenated to form *H* which is then fed into the Mamba block.

### D. Mamba Block

State Space Model (SSM) mathematically models dynamic system behavior over time by using hidden state variables that capture temporal dependencies in sequential data. SSM achieve this by defining a set of linear ordinary differential equations, which maps an input sequence to an output sequence *y*(*τ*) ∈ ℝ^*N ×N*^ at any given time interval *τ* of length *w* for *τ* ∈ [*τ*_*t*_, *τ*_*t*_ + *w*] by a latent space *s*(*τ*) ∈ ℝ^*N ×N*^ :

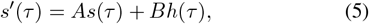

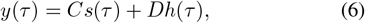

where *s*^*′*^(*τ*) is the derivative of current state *s*(*τ*), *A* ∈ ℝ^*N ×N*^ is the state transition matrix, *B* ∈ ℝ^*N ×N*^ is the input matrix and *C* ∈ ℝ^*N ×N*^ denotes the output matrix. *A* captures information about the previous state to build the new state. *B* controls how the inputs affect the state changes whereas *C* indicates how outputs are generated based on current states.

Analytically solving (5) and (6) to obtain *y*(*τ*) is challenging, as real-world data is typically discrete rather than continuous — for example, BOLD signals reflect continuous hemodynamic activity in the brain. Still, they are sampled at discrete time intervals (e.g., every 2 seconds) during fMRI acquisition. Therefore, we discretize (5) and (6) at time *t* of sampling times of fMRI time-series as:

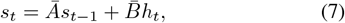

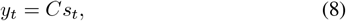

where *Ā* = exp(Δ*A*), and 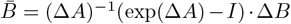. Here, Δ = [*τ*_*t*_, *τ*_*t*+1_], which represents the step size between consecutive sliding windows. Additionally, we assume *Dh*(*τ*) = 0, an exclusion justified by its analogy to a skip connection in deep learning models.

However, the main limitation of the above SSM is its lack of context awareness. To address this issue, Mamba [13] was introduced, incorporating a data-dependent selective scan mechanism to extract temporal features from dynamic FC matrices. Due to the recurrent structure employed in Mamba, each node (ROI) from the attention embeddings is updated based on the hidden states of the nodes that precede it in the sequence [22]. To ensure that Mamba prioritizes nodes with higher significance, we use the ambivert degree [23] as a heuristic to sort nodes based on their importance. The ambivert degree is a measure that captures nodes with a high intra-modular degree and the node’s participation coefficient. The rationale behind this approach is that more critical nodes should have access to more context (a more extended history of preceding nodes). By using the node ambivert degree, we identify key modular hubs and order the nodes such that Mamba prioritizes these over non-hubs. Finally, the output from the Mamba block, 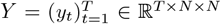, is passed through the Top-K Sliding Window Selection block.

### E. Top-K Sliding Window Selection

Deep neural networks can model complex patterns in dynamic FC. Still, they often incur high computational costs and risk overfitting since many FC features might be irrelevant in distinguishing subject groups [24]. To extract salient features from sliding windows, we propose an algorithm inspired by the STATIS method [25]. Initially designed for settings where multiple experts assess a shared set of items, it models each evaluation as a separate data matrix, enabling integrated multivariate analysis. We apply STATIS to the spatio-temporal embedding matrices to identify dominant patterns across all sliding windows and retain those that exhibit the highest similarity to the identified consensus structure. The motivation for applying this algorithm to spatio-temporal embeddings rather than directly to input-level features such as FC matrices stems from the limitation that fails to capture indirect or higher-order relationships from FC when only paired correlations are considered. Applying the top-K sliding window selection algorithm (Algorithm 1) at a later stage allows for feature selection on a richer temporal representation and improves comparability across time windows.

Given all dynamic FC matrices, we quantify their pairwise similarity using RV (Robert and Escoufier) coefficients, which is a multivariate extension of the squared Pearson correlation coefficient, that measures the similarity between two sets of points or matrices. We use the RV coefficient because it generalizes to the matrix level, capturing global rather than element-wise similarity. Its bounded range between 0 (no similarity) and 1 (identical structure) makes it highly interpretable. Dynamic FC matrices are first transformed into cross-product matrices and their inner product is then computed. Since these matrices are positive semi-definite, their inner product can be interpreted geometrically as a scalar product between two such matrices. This product is proportional to the cosine of the angle between them. This directly aligns with the RV coefficient, mathematically confirming its suitability for our algorithm. The first eigenvector of the resulting RV similarity matrix is then used to derive weights for each connectivity matrix across the sliding windows. Using these weights, we select the top-K windows, ranking them based on their contribution to the dominant connectivity patterns observed over time. Algorithm 1 provides a summary of the proposed algorithm.

#### Algorithm 1

Top K Sliding Window Selection Module

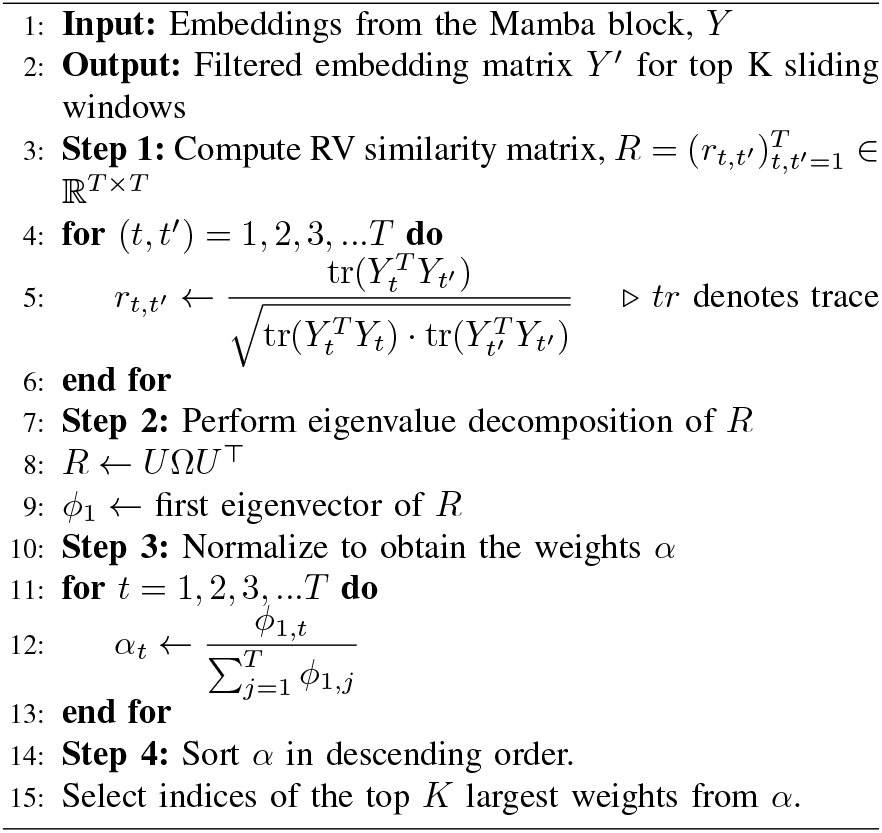

The output embeddings generated by the algorithm, 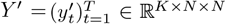, are subsequently processed using a 1D adaptive max pooling operation. The pooling operation serves to coarsen the graphs further and project the embeddings into a lower-dimensional space. The pooled embeddings are then fed into a two-layered MLP with a final layer performing softmax classification. The entire model training process is carried out in an end-to-end fashion by minimizing the cross-entropy loss function.

## III. Results

### A. Datasets and Preprocessing

ADHD-200 [26] dataset includes rs-fMRI scans from 307 individuals diagnosed with ADHD and 705 age-matched typical controls, collected from multiple sites. Data from the Athena pipeline was used for preprocessing ADHD-200. Dataset from the UCLA Consortium for Neuropsychiatric Phenomics [27] and CNI-TLC (Connectomics in NeuroImaging Transfer Learning Challenge) dataset [28] were used to validate our findings further. In the UCLA dataset, 40 out of the initial 43 participants diagnosed with ADHD were retained for analysis after preprocessing. To ensure class balance, 40 typically developing controls were randomly selected from a pool of 130. CNI-TLC comprised 100 individuals diagnosed with ADHD and 100 typically developing controls. Following recommended practices [29], we constructed dynamic FC matrices by segmenting the BOLD signals into overlapping windows, each 60 seconds in length with a 1 second stride. Both the Power Atlas [30] and the Craddock Atlas [31] were used to define 264 and 200 distinct brain regions of interest, respectively. FC matrices were computed by calculating the Pearson correlation between the mean activation time series of each ROI pair. These dynamic FC matrices, generated for each sliding window, served as inputs to our proposed model.

### B. Implementation Details

For all experiments, model training and evaluations were conducted using five different seeds. All three datasets were divided into training, validation, and test sets at a 6:2:2 ratio. Power Atlas (*N* = 264) was applied to the ADHD-200 and UCLA datasets while Craddock Atlas (*N* = 200) was used for the CNI-TLC dataset. This setup was selected because the CNI-TLC dataset was publicly available only in a preprocessed format using the Craddock Atlas, with no access to the raw fMRI data. As the Craddock Atlas is probabilistic, converting it to a deterministic atlas such as the Power Atlas [32] was not feasible. For the ADHD-200 dataset, we analyzed in a site-wise manner on NYU, OHSU, and PKU. The remaining sites were used only for biomarker analysis due to the small sample size and extreme class imbalance (NI: 20/28, KKI: 22/61, PITT: 6/95, WUSTL: 0/61 (ADHD/HC)).

The distribution of typically developing controls and ADHD subjects across datasets is as follows: UCLA – 40/40; CNI-TLC – 100/100; ADHD-200: NYU – 184/212, OHSU – 112/125, PKU – 78/116. BRAINMAP’s parameters were tuned using the validation set, and all reported results correspond to performance on the test set. Gradient descent was done using the Adam optimizer with a learning rate of 0.001. Training was carried out for 20 epochs. The hidden layer dimension in the GNN layer was set to 128. In our experiments, we used a batch size of 4, which required approximately 4.2 GB of GPU memory. All models were implemented using Python 3.9 and PyTorch 2.1 on an NVIDIA A100 GPU.

### C. Comparison Study

To assess the effectiveness of our framework, we implemented various state-of-the-art disease-specific models for dynamic FC. These models include STGCN [9], STAGIN [11], MDGL [33], DRAT [34], OT-MCSTGCN [35], Jamba [36], MSSTAN [37] and dFCExpert [38]. In STGCN, dynamic FC networks are modeled as spatio-temporal graphs to learn and capture the non-stationary patterns of FC for diagnostic tasks. STAGIN utilizes GIN to extract graph-level representations of the brain connectome at each time step, and employs a transformer encoder to embed temporal information across the dynamic graph sequence. MDGL constructs multi-scale dynamic brain graphs by utilizing two distinct brain atlases to improve disorder prediction. DRAT fuses static and dynamic FC features through multi-level attention mechanisms, thereby enhancing representation learning. OT-MCSTGCN leverages OT to model the evolving topology of dynamic brain networks and employs a multi-channel spatio-temporal GCN to jointly extract temporal and spatial features from fMRI and DTI modalities. Jamba integrates Transformer and Mamba blocks to effectively address long-context natural language processing tasks. MSSTAN is based on a topology-enhanced Graph Transformer that captures long-term temporal dependencies in functional connectomes across multiple scales. Lastly, dFCExpert models the brain modularity and learns robust representations of dynamic FC patterns in fMRI data using a GNN and Mixture of Experts (MoE) framework. Default model parameters were used to train all the models.

We found that our proposed model outperformed existing models in all 40 experiments (with statistically significant improvements in 27 experiments based on Student’s t-test, and among these, 7 showed medium-to-large effect sizes according to Cohen’s d-test) across all three datasets. Detailed model performances are presented in Table I^1^. Our approach achieves superior performance by efficiently modeling spatio-temporal dynamics via the integration of high-level distributional and graph representations in dynamically evolving brain networks. Combining attention’s capacity for short-range dependency modeling with Mamba’s efficiency in capturing long-range patterns enhances prediction accuracy. The Top-K Sliding Window Selection complements this by reducing overfitting, thereby enhancing generalizability over dynamic FC models that rely on the entire feature space. Overall, these results demonstrate the superior classification performance of our proposed model.

**TABLE I.**
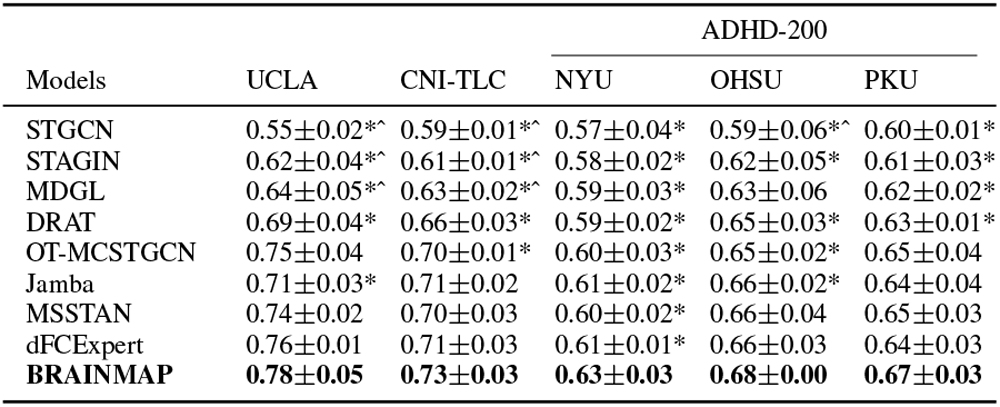
Comparison with the ADHD prediction accuracies of existing models.

### D. Ablation Study

To validate the effectiveness of our proposed method, we conducted ablation studies on all three datasets. The experimental results, summarized in Table II^1^, demonstrate a significant decline in performance when the Mamba block and Cross-Attention layer (Attn) are omitted. This highlights their crucial role in effectively filtering highly predictive signals across all time points. Furthermore, removing the Top-K selection algorithm led to a noticeable performance drop, suggesting that specific sliding windows lead to better generalization and play a crucial role in identifying more reliable biomarkers. A core strength of BRAINMAP lies in its dual-representation strategy, which captures both the global distribution and spatial characteristics of dynamic FC. Eliminating these representations resulted in an inferior performance compared to the complete architecture. Our ablation experiments show the utility of BRAINMAP in distinguishing between typically developing controls and ADHD subjects.

**TABLE II.** Ablation study of BRAINMAP: ADHD prediction accuracies.

| Models | UCLA | CNI-TLC | ADHD-200 |  |  |
| --- | --- | --- | --- | --- | --- |
|  |  |  | NYU | OHSU | PKU |
| w/o Top-K | 0.71±0.03* | 0.70±0.03 | 0.60±0.01* | 0.63±0.05* | 0.62±0.04* |
| w/o Mamba | 0.65±0.03*^ | 0.68±0.02* | 0.57±0.01* | 0.59±0.04*^ | 0.58±0.01*^ |
| w/o Attn | 0.69±0.04* | 0.66±0.02* | 0.58±0.01* | 0.62±0.03* | 0.61±0.02* |
| w/o OT | 0.71±0.05 | 0.72±0.02 | 0.59±0.01 | 0.61±0.03*^ | 0.60±0.01* |
| w/o GNN | 0.74±0.02 | 0.71±0.01 | 0.61±0.02 | 0.64±0.03* | 0.64±0.02 |
| **BRAINMAP** | **0.78±0.05** | **0.73±0.03** | **0.63±0.03** | **0.68±0.00** | **0.67±0.03** |

### E. Performance across different GNNs

In Table III^1^, we further show how the performance of the BRAINMAP framework varies when integrated with different GNN architectures. Specifically, we evaluate the GCN model [39], ChebGCN (Cheb) [40], GraphSAGE (SAGE), GAT [41], GraphGPS (GrGPS) [42], GraphFormer (GrForm) [43], and GIN. Among these, GIN, shown to be as powerful as the Weisfeiler-Lehman (WL) graph isomorphism test, achieves superior performance across the three datasets. This aligns with its established advantage in effectively capturing complex structural patterns in graph classification tasks. Based on these results, we adopted the GIN architecture as the default GNN backbone in BRAINMAP for all downstream analyses.

**TABLE III.** Comparison of prediction accuracies with different GNNs in BRAINMAP.

| GNNs | UCLA | CNI-TLC | ADHD-200 |  |  |
| --- | --- | --- | --- | --- | --- |
|  |  |  | NYU | OHSU | PKU |
| GCN | 0.66±0.03*^ | 0.67±0.03* | 0.59±0.02* | 0.64±0.02* | 0.62±0.02* |
| Cheb | 0.61±0.02*^ | 0.66±0.01* | 0.57±0.02* | 0.63±0.02*^ | 0.60±0.02* |
| SAGE | 0.71±0.03* | 0.69±0.02* | 0.61±0.01* | 0.66±0.02 | 0.64±0.03 |
| GAT | 0.74±0.02 | 0.72±0.01 | 0.59±0.02* | 0.65±0.01* | 0.64±0.01* |
| GrGPS | 0.74±0.03 | **0.73±0.03** | **0.64±0.02** | 0.67±0.02 | 0.65±0.01 |
| GrForm | 0.75±0.04 | 0.72±0.00 | **0.64±0.02** | 0.66±0.03 | 0.64±0.03 |
| **GIN** | **0.78±0.05** | **0.73±0.03** | 0.63±0.03 | **0.68±0.00** | **0.67±0.03** |

### F. Performance across node ordering metrics

Table IV^1^ summarizes the classification performance of BRAINMAP using various node ordering metrics within the Mamba block. We compare the ambivert degree (Amb. Degree) with modular degree (Mod. Degree) [44], gateway coefficient (Gate Coeff) [45], and a baseline scenario with no node ordering applied. The node ordering strategy yielded a significant performance improvement on all three datasets compared to the baseline, where no ordering was used within the Mamba block. Specifically, the ambivert degree consistently outperforms the other metrics, highlighting that ordering nodes based on hub prominence and modular properties enables the Mamba block to capture temporal features more effectively. Thus, the ambivert degree was selected as the primary node ordering metric for all subsequent analyses.

**TABLE IV.**
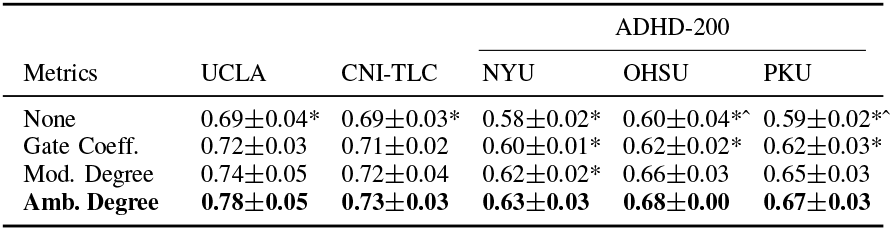
Comparison of prediction accuracies with different node ordering metrics.

### G. Evaluation of BRAINMAP for different K values

Table V^1^ reports the predictive performance of BRAINMAP on the three datasets under varying values of K in the Top-K Sliding Window Selection algorithm, with K set to 30, 40, 50, 60, 70. The results indicate that K=50 yields strong performance across most settings, except for site OHSU from the ADHD-200 dataset. A general upward trend in performance is observed as K increases from 30 to 50, followed by a decline as K approaches 70. This analysis highlights the model’s sensitivity to the choice of K and its impact on classification performance. Based on this empirical evaluation, we select K = 50 as the default value for downstream analyses.

**TABLE V.**
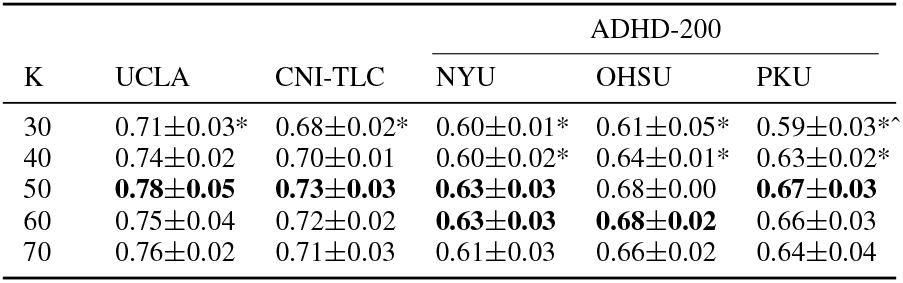
comparison with different values for k in top-k selection.

### H. Leave-One-Site-Out Cross-Validation

To assess the reliability of BRAINMAP, we performed leave-one-site-out cross-validation on the ADHD-200 dataset. This serves as an external validation and accounts for site-specific effects. For this analysis, we focused on four sites: NI, NYU, OHSU, and PKU. The remaining sites were excluded due to extreme class imbalance. In each case, we trained the model on data from three sites (e.g., NYU, OHSU, and PKU) and tested it on the held-out site (e.g., NI), and repeated this process for all sites. From Table VI^1^, it is evident that BRAINMAP consistently outperforms state-of-the-art models in cross-validation performance. This suggests that the model captures invariant features of ADHD rather than relying on site-specific signatures. Thus, BRAINMAP achieves strong performance across both individual site analyses and cross-validation, underscoring its reliability and strong generalizability across multiple sites and datasets.

**TABLE VI.** Comparison study of BRAINMAP with Leave-One-Site-Out Cross-Validation on ADHD-200.

| Models | NI | NYU | OHSU | PKU |
| --- | --- | --- | --- | --- |
| STGCN | 0.50±0.03*<sup>^</sup> | 0.51±0.01*<sup>^</sup> | 0.53±0.01*<sup>^</sup> | 0.52±0.01*<sup>^</sup> |
| STAGIN | 0.54±0.05*<sup>^</sup> | 0.53±0.02* | 0.53±0.02*<sup>^</sup> | 0.53±0.02*<sup>^</sup> |
| MDGL | 0.58±0.02*<sup>^</sup> | 0.54±0.01* | 0.55±0.01* | 0.55±0.02* |
| DRAT | 0.60±0.01*<sup>^</sup> | 0.56±0.03 | 0.56±0.02* | 0.57±0.02* |
| OT-MCSTGCN | 0.65±0.03 | 0.57±0.03 | 0.58±0.03 | 0.58±0.02 |
| Jamba | 0.64±0.02* | 0.57±0.01 | 0.59±0.01 | 0.57±0.03 |
| MSSTAN | 0.65±0.02 | 0.55±0.04 | 0.57±0.02 | 0.59±0.03 |
| dFCExpert | 0.66±0.03 | 0.57±0.02 | 0.60±0.01 | 0.60±0.02 |
| **BRAINMAP** | **0.68±0.02** | **0.58±0.02** | **0.61±0.02** | **0.61±0.03** |

## IV. Biomarker Analysis

Diagnostic biomarkers generated by state-of-the-art explainable AI techniques exhibit significant variation across explainers and predictors [46]. This variability underscores the need for cautious interpretation of top-ranked features with high saliency scores – features that are often emphasized in existing studies. Consequently, the common practice of validating potential biomarkers by merely referencing prior literature that reports similar findings may be insufficient to establish their actual relevance. We employ quantitative evaluation metrics introduced in the RE-CONFIRM framework, under the hypothesis that our model may yield robust biomarkers and generalize reasonably across datasets and explainers. The biomarker analysis is done using RE-CONFIRM: a framework designed to evaluate the robustness of salient FC features identified by classification models and explanation methods, which comprises nine distinct metrics designed to assess core properties of explanation methods and metrics, specifically within the context of brain connectomes.

Decoding of potential biomarkers was performed using BRAINMAP under four conditions: (i) on individual sites within the ADHD-200 dataset; (ii) on the entire ADHD-200, UCLA, and CNI-TLC datasets without ComBat harmonization [47]; (iii) on the same combined datasets with ComBat harmonization; (iv) on each ADHD subtype within the ADHD-200 dataset. To account for site-related variability, such as scanner differences and pre-processing pipelines, when combining datasets, we applied ComBat as a data harmonization technique to mitigate batch effects. Each chord diagram in Figures 3, 4, and 5 represents the 25 most salient connections (corresponding to the top 0.1% of features). Saliency scores in BRAINMAP were derived using attention weights from the Cross-Attention layer, aggregated across all five seeds. Scores were computed for the Top-K segments. These values were then averaged and standardized. The analysis was carried out at the modular level.

**Fig. 3.**
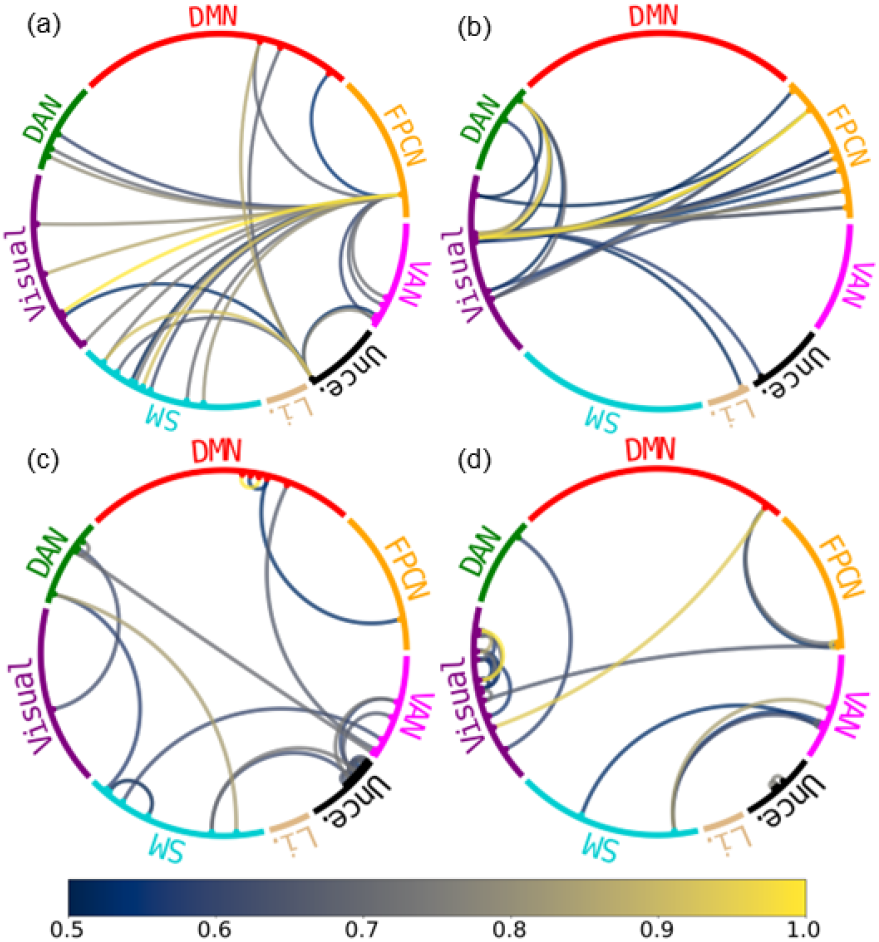
Chord diagrams representing site-invariant salient FC connections for UCLA, CNI-TLC, ADHD-200 with ComBat, and all ADHD datasets with ComBat. In each chord diagram, only the top 25 connections were visualized to reduce cluttering. (a) UCLA (b) CNI-TLC (c) Whole ADHD-200 with ComBat (d) All ADHD datasets with ComBat. Brain network labels as follows:-DMN: Default Mode Network; FPCN: Frontoparietal control network; VAN: Ventral Attention Network; Unce.: Uncertain; Li.: Limbic Network; SM: Sensorimotor Network; Visual: Visual network; DAN: Dorsal Attention Network.

**Fig. 4.**
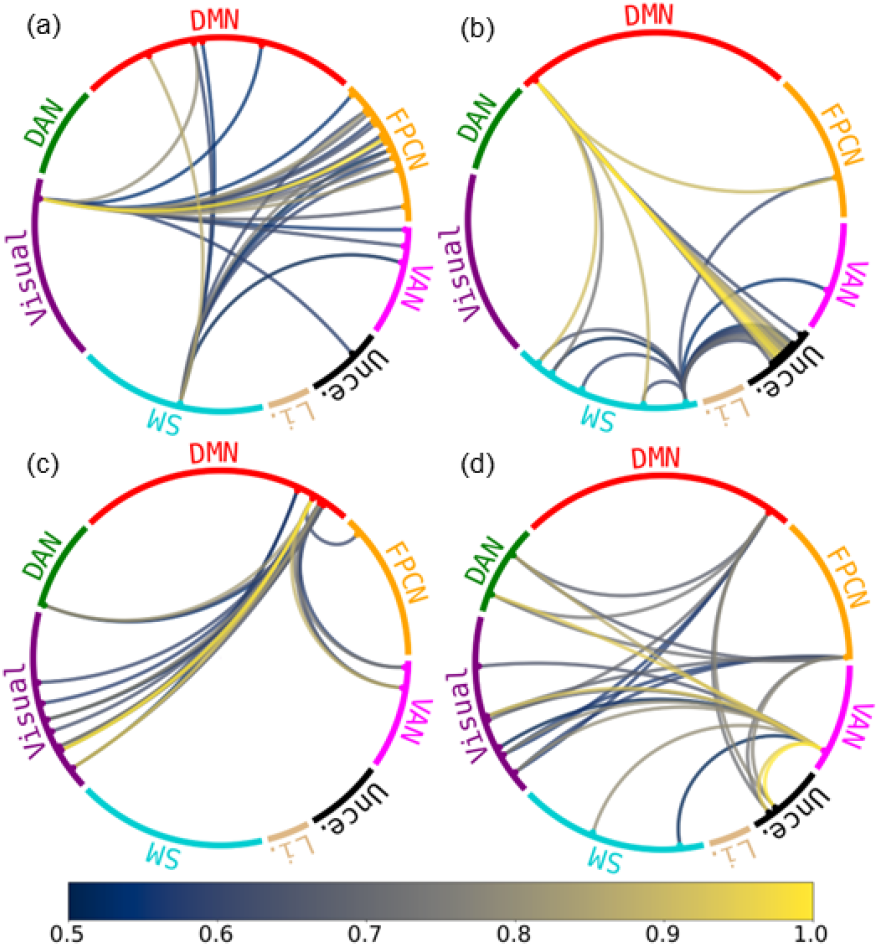
Chord diagrams representing site-invariant and site-wise salient FC connections of the ADHD-200 dataset. In each chord diagram, only the top 25 connections were visualized to reduce cluttering. (a) Whole ADHD-200 (b) NYU (c) OHSU (d) PKU. Brain network labels as follows:-DMN: Default Mode Network; FPCN: Frontoparietal control network; VAN: Ventral Attention Network; Unce.: Uncertain; Li.: Limbic Network; SM: Sensorimotor Network; Visual: Visual network; DAN: Dorsal Attention Network.

**Fig. 5.**
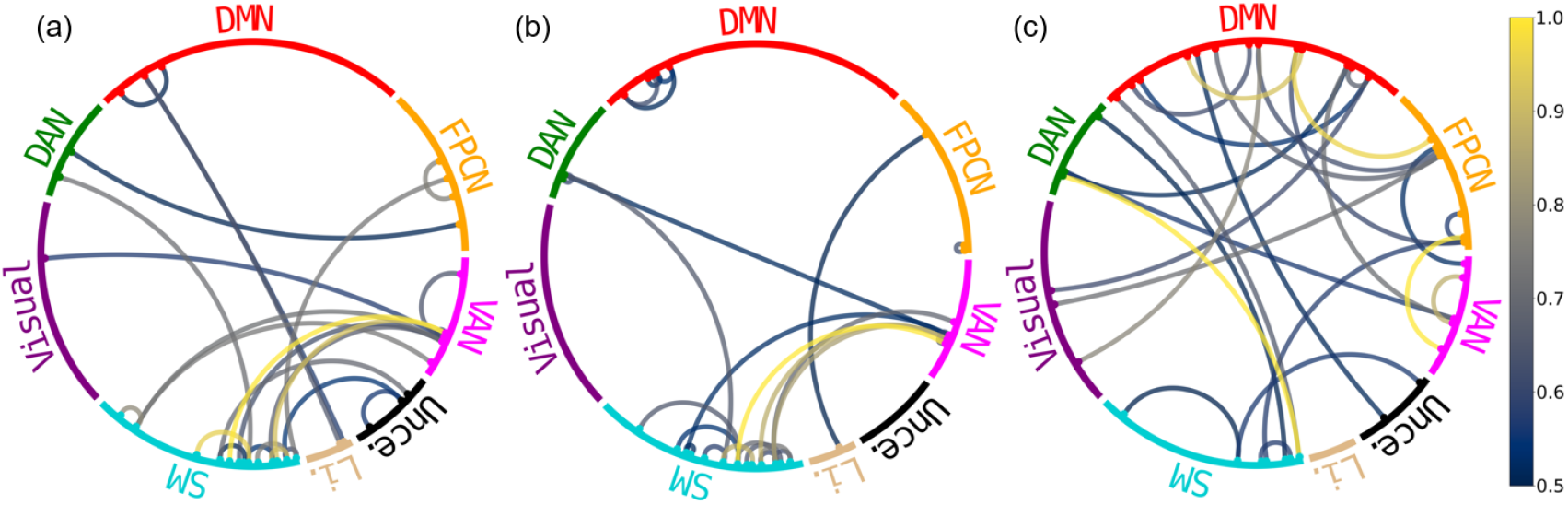
Chord diagrams representing subtype-specific salient FC connections of the ADHD-200 dataset. In each chord diagram, only the top 25 connections were visualized to reduce cluttering. (a) ADHD-combined (b) ADHD-hyperactive/impulsive (c) ADHD-inattentive. Brain network labels as follows:-DMN: Default Mode Network; FPCN: Frontoparietal control network; VAN: Ventral Attention Network; Unce.: Uncertain; Li.: Limbic Network; SM: Sensorimotor Network; Visual: Visual network; DAN: Dorsal Attention Network

### A. Evaluating robustness of salient features

We apply RE-CONFIRM to evaluate three models: BRAIN-MAP, OT-MCSTGCN, and Jamba. The latter two models were selected due to their comparable classification performance relative to BRAINMAP. Table VII^1^ presents a summary of the RE-CONFIRM evaluation metrics for all models across the combined ADHD-200 dataset, using attention weights from the Cross-Attention layer as the explainer.

**TABLE VII.** evaluation metrics for attention scores on adhd-200.

| Metrics | OT-MCSTGCN | Jamba | BRAINMAP |
| --- | --- | --- | --- |
| MPRC (↑) | $0.52 \pm 0.03^*$ | $0.55 \pm 0.02^*$ | **$0.61 \pm 0.02$** |
| DRC (↑) | $0.42 \pm 0.03^*$ | $0.46 \pm 0.04$ | **$0.48 \pm 0.02$** |
| Fidelity+ (↑) | $0.41 \pm 0.01^*$ | **$0.47 \pm 0.02$** | $0.38 \pm 0.02^*$ |
| Stability (↑) | **$1.00 \pm 0.00$** | **$1.00 \pm 0.00$** | **$1.00 \pm 0.00$** |
| HAC (↑) | $0.32 \pm 0.01^*$ | **$0.37 \pm 0.03$** | $0.35 \pm 0.03$ |
| Fidelity- (↓) | $0.27 \pm 0.02^*$ | **$0.24 \pm 0.01$** | $0.28 \pm 0.03^*$ |
| Sens (↓) | **$0.21 \pm 0.01$** | $0.23 \pm 0.03$ | $0.23 \pm 0.01^*$ |
| II (↓) | **$0.11 \pm 0.01$** | $0.13 \pm 0.02$ | $0.17 \pm 0.01^*$ |
| MR (↓) | $0.44 \pm 0.02$ | $0.48 \pm 0.02^*$ | **$0.43 \pm 0.03$** |
| MR-20 (↓) | $0.42 \pm 0.01^*$ | $0.44 \pm 0.02^*$ | **$0.40 \pm 0.01$** |

Our results suggest that BRAINMAP performs slightly better than the other two models. It demonstrates greater stability, lower MR and MR-20 scores. This implies the salient features in BRAINMAP may exhibit a more pronounced modular relationship compared to the other two models. Models that incorporated attention mechanisms along with SSM modules in their architecture (BRAINMAP and Jamba) were found to generally perform better than models with only attention: they exhibited higher MPRC, DRC, and HAC scores. Explanations generated by Jamba capture the true behavior of the black-box model more accurately compared to those produced by other evaluated methods. Conversely, OT-MCSTGCN produces salient features that are less sensitive to parametrization changes and consistent across different seed initializations. Overall, these results suggest that BRAINMAP may be a relatively more robust predictive model compared to OT-MCSTGCN and Jamba, with its identified biomarkers demonstrating moderately greater reliability.

### B. Site-wise salient FC features

At the site NYU, biomarker analysis revealed prominent inter-modular connectivity between the Default Mode Network (DMN) and both the Sensorimotor Network (SM) and the Frontoparietal Control Network (FPCN). In addition, salient intra-modular connections within the SM were observed, indicating potential localized motor network disruptions, commonly associated with ADHD [48]. Site OHSU exhibited a biomarker profile dominated by inter-modular connections centered on the DMN. Prominent connections included the DMN–Visual Network, the DMN–Dorsal Attention Network (DAN), and the DMN–FPCN, with the latter overlapping with findings from the site NYU. In contrast, biomarker analysis at the PKU site revealed inter-modular connections that were concentrated in three areas: DAN, DMN, and Ventral Attention Network (VAN). Notably, DMN–Visual, DMN–DAN, and SM–VAN connections were also observed at the sites OHSU and NYU, respectively. When comparing salient connections across all sites, DMN, known for its attention and cognitive roles within the brain, consistently emerged as a common module of interest, aligning with previous ADHD studies [49].

### C. Site-invariant salient FC features

In the UCLA dataset, the chord diagram revealed widespread salient inter-modular connections, mainly involving the FPCN and Visual Network. Similarly, in the CNI-TLC dataset, a high density of salient connections was observed between the FPCN and Visual Network, as well as between the FPCN and DAN. This differed markedly from the observations in the ADHD-200 dataset, where both inter- and intra-modular connections involving the Visual Network were prominent and salient. Notably, the salient connection between DAN and the Visual Network was present in both the ADHD-200 and the CNI-TLC datasets, but was absent in the UCLA dataset. This may be attributed to differences in the age groups targeted across datasets: ADHD-200 and CNI-TLC primarily included children and adolescents, whereas the UCLA dataset comprised adult participants. The presence of DAN-Visual Network connection in younger cohorts suggests it may be an FC feature specific to pediatric ADHD. This observation aligns with existing literature on ADHD biomarkers in pediatric populations [50].

The DAN–Visual Network connection was again consistently observed in both the whole ADHD-200 and the combined datasets when ComBat was applied. Based on the individual dataset analysis above, this consistency likely reflects the higher proportion of children and adolescents compared to adults in the pooled age groups. Additionally, numerous intra-modular connections were also present in the pooled datasets under both conditions, with and without ComBat. The dominance of intra-modular connections suggests that dysregulation within individual brain networks, rather than disruptions between different networks, may be an important marker of ADHD across diverse populations and acquisition sites. Moreover, the continued prominence of the DAN-Visual Network connection following ComBat suggests that this salient connection is unlikely to be solely due to site-specific variability and may instead reflect underlying neurobiological factors, supporting its potential as a generalizable biomarker for ADHD.

For additional comparison, we also identified salient connections in the ADHD-200 dataset using Jamba, as it was the second-best performing model based on the RE-CONFIRM evaluation metrics. Interestingly, the DMN-SM connection emerged as the most salient. However, this connection was not prominent across other datasets or individual sites, except for the site NYU. This suggests that the biomarkers identified by other models may have limited reproducibility across most sites and datasets, indicating lower robustness. In contrast, the relative consistency of our model’s findings supports the reliability of our biomarker decoding approach.

### D. Subtype-specific salient FC features

We used data from sites NYU, OHSU, PKU, NI, and KKI of the ADHD-200 dataset for subtype-specific analysis. These were the only sites that included all three ADHD subtypes: combined (159 subjects), hyperactive/impulsive (11 subjects), and inattentive (110 subjects). The resulting chord diagrams revealed a biomarker profile largely shaped by intra-modular connections across all subtypes, with prominent involvement of SM in combined (Figure 5(a)) and hyperactive subtypes (Figure 5(b)) and DMN in the inattentive subtype (Figure 5(c)). We also found that the connection between VAN-SM is most salient for both combined and hyperactive subtype. Notably, it is not present in the inattentive subtype. Salient connections at inattentive subtype included DMN-FPCN, DAN-SM and FPCN-VAN. This suggests that VAN-SM connection could be an FC feature specific to the hyperactive subtype. While the combined and hyperactive subtypes have distinct salient connections, the involvement of VAN and SM, known for their role in stimulus-driven attention and motor control within the brain, aligns with findings from prior ADHD studies [51].

We also observed a statistically significant similarity between the combined and hyperactive subtypes (Pearson’s r = 0.71, p *<* 0.05). Correlations between combined and inattentive subtypes were moderate (r = 0.54), correlation between inattentive and hyperactive subtypes was slightly weaker (r = 0.44). Overall, our results suggest that the VAN-SM pattern may reflect a stronger influence of hyperactive traits within the combined subtype. This is particularly noteworthy given the small sample size of the hyperactive subtype.

### E. Association study with Conners Scale

To further substantiate the identified salient functional connections, their FC values were correlated with symptom severity measures of the disorder, such as the Conners Scale in the case of ADHD. The Conners Scale is a behavioral rating tool used to assess ADHD symptoms and related issues in children and adolescents, based on parent, teacher, or self-reports, using standardized T-scores. For our analysis, we selected 100 subjects each from sites NYU and PKU within the ADHD-200 dataset. These two sites were the only ones that provided Conners Scale (Parent Rating) metadata, making them suitable for our study.

To examine these associations, FC values of the top salient modular connection were correlated with Conners Scale scores. FC values were derived by first grouping 264 Power Atlas ROIs according to their module assignments and averaging the time series within each module. An FC matrix was then constructed by computing the Pearson correlation between these modular time series. For this analysis, we chose the most salient sliding window, as identified by our Top-K Sliding Window Selection algorithm. FC matrices exhibited values ranging from min = –0.999 to max = 1.000, with mean of 0.018 and a standard deviation of 0.299. We observed a positive correlation between the DAN–Visual Network connection and Conners Scale scores. This relationship is visualized in Figure 6(a) using scatter plots of FC values against the Conners Scale. Given the small sample size, we evaluated the stability of the FC values–Conners score association using nonparametric bootstrap resampling (10,000 iterations). This analysis yielded a consistent association (*ρ* = 0.691) with a 95% confidence interval of [0.236, 0.893]. Although the interval was broad due to the limited sample size, the bootstrap mean estimate showed minimal shrinkage relative to the observed value (Δ*ρ* = 0.021), indicating low sampling bias and supporting the robustness of the observed relationship.

**Fig. 6.**
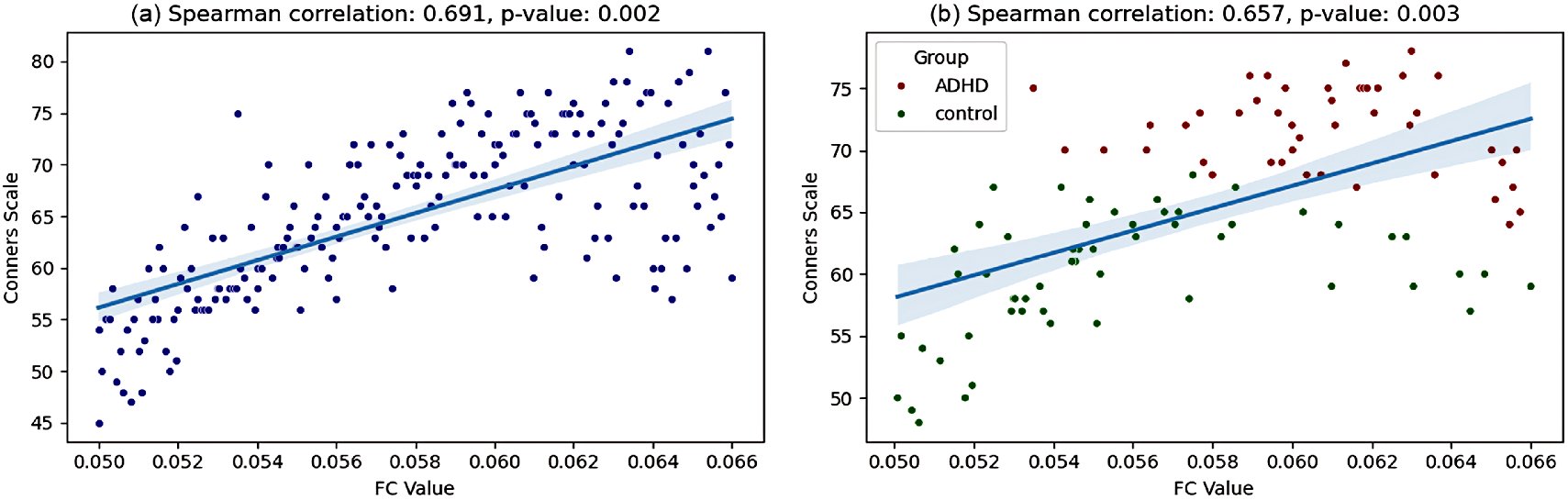
Scatter plots with regression lines showing the relationship between FC values of the DAN–Visual Network connection and Conners Scale scores: (a) includes data from both sites NYU and PKU, while (b) contains data from only site PKU.

Additionally, we examine this association at site PKU in Figure 6(b) by color-coding the data points according to the diagnostic labels. Typically developing controls primarily exhibit lower FC values and Conner Scale scores, whereas individuals with ADHD typically show higher values on both measures. These findings suggest the potential relevance of the identified biomarkers, as the FC changes identified by our model are associated with the severity of ADHD symptoms. One notable limitation of this analysis is the small sample size. Therefore, this serves solely as a proof of concept. Further studies utilizing larger fMRI datasets that include Conners Scale scores are necessary to validate these associations.

## V. Discussion

Several key findings emerge from this study. Firstly, BRAINMAP shows strong performance on three public ADHD datasets, surpassing six state-of-the-art dynamic FC models. This can be attributed to BRAINMAP’s ability to leverage the complementary strengths of attention mechanism, Mamba block, and OT architectures, effectively capturing both short-term and long-range dependencies across sliding windows. Another key factor contributing to Mamba’s superior performance is the node ordering strategy, where each node is updated based on the hidden states of preceding nodes in the sequence. Ordering nodes by ambivert degree introduces a biologically informed sequence that emphasizes hubs and connector nodes, enabling the model to capture hierarchical dependencies and hub-driven interactions. This inductive bias allows BRAINMAP’s Mamba block to model temporal dynamics more effectively while preserving meaningful network topology. And unlike Transformers, Mamba excels at capturing complex long-range dependencies in sequential data [52]. Our experiments using the Top-K Selection algorithm revealed that the proposed model not only reduces overfitting but also shows a degree of redundancy within the sliding window features. BRAINMAP achieves its highest predictive accuracy when utilizing only the top 50 most salient sliding windows. The integration of OT and GNNs consistently enhanced performance compared to when they were excluded, supporting our initial hypothesis that correcting distributional shifts and capturing spatial features leads to improved generalization and classification performance. Specifically, GIN proved to be the most effective within the BRAINMAP framework, outperforming other GNNs in our evaluations. This aligns with existing works, which indicate that GIN outperforms other GNN architectures in capturing complex graph structures, making it effective for graph classification tasks [53]. BRAINMAP also achieved strong classification performance in leave-one-site-out cross-validation on ADHD-200 dataset, with performance trends consistent with those observed in the site-wise analyses. These results provide strong evidence of its generalizability across multiple sites and datasets. We further examined BRAINMAP’s calibration and fairness with respect to demographic factors such as age, sex, and ADHD subtype. These factors had minimal and inconsistent impact on the predictive performance, demonstrating BRAINMAP’s stability to demographic variability. Notably, the slight variations were observed in younger cohorts than adult participants.

Our method identified site-wise salient FC features, as well as site-invariant and subtype-specific biomarkers. We observed that the majority of salient connections involved the DMN, which is well-documented to be disrupted in individuals with ADHD. Notably, across both the ADHD-200 and CNI-TLC datasets, BRAINMAP highlighted the connection between the DAN and the Visual Network as salient. This connection was also observed when all sites within ADHD-200 and all three datasets were combined and harmonized using ComBat. Additionally, intra-modular connections were prominent in the harmonized setting in both cases and across various subtypes. This highlights their potential role in ADHD-related disruptions across diverse populations and age groups. The recurrence of the salient DAN–Visual Network connection across site-wise, site-invariant, and data harmonization analyses suggests that it is a prominent biomarker identified by BRAINMAP. This biomarker has also been reported in other disorders associated with attentional difficulties [54]. Our subtype-specific analysis revealed that the VAN–SM connection may be an FC feature specific to the hyperactive subtype. It was present in both hyperactive and combined subtypes but absent in the inattentive subtype. These findings suggest that different analytical conditions may produce varying sets of biomarkers. This indicates that methodological choices can influence how ADHD heterogeneity is represented and the identification of subtype or population-specific neural signatures.

Furthermore, we employed the RE-CONFIRM framework to validate the robustness of the salient features identified by our model. BRAINMAP generally performed better than OT-MCSTGCN and Jamba on most RE-CONFIRM metrics, supporting the relative reliability of our biomarker analysis. We also demonstrated the practical relevance of the identified biomarkers by examining the associations between ADHD symptom severity and their corresponding FC values. A positive correlation was found between FC values of the DAN–Visual Network connection and Conners Scale scores. By linking FC changes with symptom trajectories, these associations suggest that our biomarkers may have potential for monitoring treatment efficacy and disease progression in clinical trials. Through our analysis, we highlight reproducible FC patterns that may contribute to understanding ADHD and its symptom severity, without claiming immediate clinical diagnostic utility.

One limitation of this study is that the optimality of the Top-K parameter in the sliding window selection algorithm is empirically determined and may not generalize to other disorders or acquisition settings. Future work could investigate how this parameter varies across different neuro-developmental disorders and site-related parameters. Another limitation is that our classification and biomarker analysis were conducted in a site-wise manner, without extending to multi-site analyses. Pooling datasets from multiple sites for similar disorders requires careful consideration of several challenges, including site-related variability, inconsistent labeling practices, and, most importantly, concerns about data privacy. Future work could focus on extending our approach to a federated learning setting, particularly within the dynamic FC setting, enabling decentralized model training without the need for direct data sharing. Identification of specific imaging biomarkers for ADHD is made more challenging by the presence of ADHD subtypes [55]. One potential direction for future work is to investigate subtype-specific differences in FC for ADHD using a larger dataset. Finally, although our method has been evaluated only on ADHD neuroimaging datasets, it applies to other neurodevelopmental and psychiatric conditions, such as Autism and Schizophrenia.

## VI. Conclusion

We presented BRAINMAP, a framework that leveraged the combined strengths of attention mechanisms, Mamba blocks, and representations derived from Optimal Transport and Graph Neural Networks within a dynamic FC setting. We introduced a Top-K sliding window selection algorithm designed to reduce overfitting in dynamic FC models. We show that our model outperforms previous state-of-the-art models based on dynamic FC. We conduct extensive experiments to evaluate the effectiveness of our model on three publicly available ADHD datasets: ADHD-200, UCLA, and CNI-TLC. Our biomarker analysis revealed the DAN–Visual Network connection as a salient FC feature specific to ADHD, while most site-wise connections were predominantly centered around the DMN. Ultimately, our model identifies biomarkers that are potentially robust and clinically relevant, with possible implications for improving the understanding and assessment of ADHD.

## Footnotes

1 * indicates statistical significance (p-value *<* 0.05), ^ indicates medium-to-large effect size (d *>* 0.5).

---

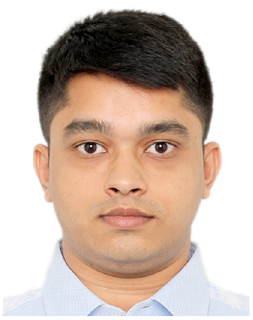

**Deepank Girish** holds a Bachelor’s degree in Computer Science and Engineering from PES University, Bengaluru, India. He is currently pursuing a Ph.D. in the College of Computing and Data Science at Nanyang Technological University (NTU), Singapore. His research focuses on applying deep learning techniques to functional MRI data, with a particular emphasis on dynamic functional connectivity. His work primarily involves identifying robust biomarkers for neurodevelopmental disorders using explainable AI approaches. Additionally, he explores the application of graph neural networks and sequence-based models to develop efficient and accurate disease detection architectures.

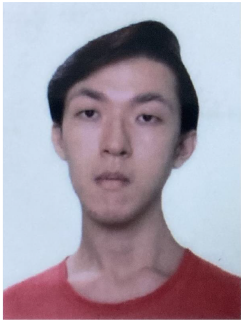

**Yi Hao Chan** is a Postdoctoral Fellow at the College of Computing and Data Science, Nanyang Technological University (NTU), Singapore. He earned both his undergraduate degrees, specializing in Business Analytics, Data Science, and Artificial Intelligence, as well as his Ph.D. in Computer Science from NTU. His doctoral research focused on brain imaging, multimodal learning, and explainable AI. His current research lies at the intersection of Machine Learning and Neuroscience, with a particular emphasis on temporal graph interpretability.

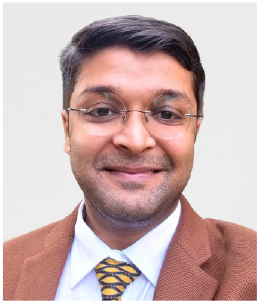

**Sukrit Gupta** is an Assistant Professor in the Department of Biomedical Engineering and the School of Artificial Intelligence and Data Engineering at the Indian Institute of Technology (IIT) Ropar, India. Before joining IIT Ropar, he was a Research Fellow at the Hasso Plattner Institute in Berlin and continued as an Adjunct Fellow. He also worked as a Research Scientist at ASTAR, Singapore, and later as a Visiting Scientist. His current research lies at the intersection of Artificial Intelligence and Healthcare, where he develops computational models to analyze diverse biological and medical datasets.

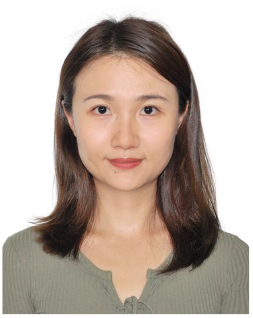

**Jing Xia** is a Hundred Talents Program Researcher at the College of Instrument and Computer Science, Zhejiang University. She received her B.S. and D.E. degrees in the Department of Computer Science and Technology at Shandong University. She conducted postdoctoral research at the College of Computing and Data Science, Nanyang Technological University, Singapore, and the School of Biomedical Engineering, National University of Singapore. Her research interests include brain functional connectivity, deep learning, and medical image analysis.

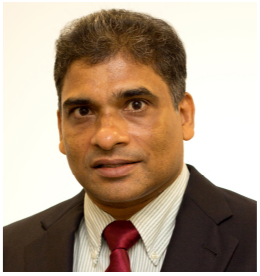

**Jagath C. Rajapakse** (Fellow, IEEE) is a Professor at the College of Computing and Data Science, Nanyang Technological University (NTU), Singapore. His current research focuses on developing advanced computational methods and tools for diagnosing and treating brain disorders by integrating neuroimaging with multi-omics data. His research interests are in explainable AI, generative AI, brain imaging, and computational and systems biology. He is a Fellow of IEEE and currently serves as Editor for Engineering Applications of Artificial Intelligence. He has previously served as Associate Editor for IEEE Transactions on Medical Imaging, IEEE Transactions on Neural Networks and Learning Systems, and IEEE Transactions on Computational Biology and Bioinformatics.

